# The irregular fruit green netting: An eggplant domestication trait controlled by the *SmGLK2* gene with implications in fruit colour diversification

**DOI:** 10.1101/2023.06.28.546667

**Authors:** Andrea Arrones, Silvia Manrique, Virginia Baraja-Fonseca, Mariola Plazas, Jaime Prohens, Ezio Portis, Lorenzo Barchi, Giovanni Giuliano, Pietro Gramazio, Santiago Vilanova

## Abstract

The distribution of chlorophylls in the eggplant (*Solanum melongena*) fruit peel can be uniform or display an irregular green netting pattern. The fruit green netting phenotype, manifested as a gradient of dark green netting, more intense in the proximal part of the fruit on a pale green background, is commonly present in eggplant wild relatives as well as in some eggplant landraces. During domestication and modern breeding of eggplant, uniform fruit colour has been selected. However, the fruit green netting contributes to a greater diversity of fruit colours. Here, we have used over 2,300 individuals from several germplasm and experimental populations, including a multi-parental MAGIC population for candidate genomic region identification, an F_2_ population for BSA-Seq, and advanced backcrosses for edges-to-core fine mapping, to determine that *SmGLK2* is the gene underlying the irregular netting in eggplant fruits. We have also analysed the gene sequence of 178 *S. melongena* accessions and 22 wild relative species for tracing the evolutionary changes that the gene has undergone over the course of domestication. Three different mutations were identified leading to the absence of netting. The main causative indel results in the appearance of a premature stop codon disrupting the protein conformation and function, which was confirmed by Western blotting analysis and confocal microscopy observations. *SmGLK2* has a major role in the regulation of chlorophyll biosynthesis in eggplant fruit peel, and therefore in eggplant fruit photosynthesis.

## Introduction

Variation in the presence or absence of chlorophylls in the eggplant (*Solanum melongena* L.) fruit peel contributes to the wide range of fruit colours in this species (Arrones *et al*., 2022). Furthermore, the distribution of chlorophylls can be uniform or irregular, the latter being referred to as netting, reticulation, or variegation (Tigchelaar, 1968; Daunay *et al*., 2004). The presence of the fruit green netting (FN) trait is commonly present in eggplant wild species and landraces. Cultivated eggplant differs morphologically and physiologically from its wild ancestors. Indeed, wild eggplants produce small, bitter, green-netted fruits, while domesticated eggplants are characterized by desirable agricultural traits which were directly and indirectly selected and fixed during domestication (Frary and Doğanlar, 2003; Fuller, 2007). These include big and non-bitter fruits, with a diversification of immature fruit peel colour, resulting from the combination of chlorophyll and anthocyanin pigments (Taher *et al*., 2017; Page *et al*., 2019; Page *et al*., 2021). While numerous studies focus solely on the presence or absence of chlorophylls in the eggplant fruit peel, it is noteworthy that reticulated fruits often exhibit a gradient in netting density in a pale green background. This gradient often leads to a deep green colour in the proximal portion of the fruit, and gradually diminishes towards the distal end. Thus, FN contributes to further increase the colour diversity of eggplant, resulting in an irregular green or dark purple colour depending on the concurrent absence or presence of anthocyanins.

The gradient of expression of this trait is reminiscent of the green shoulder trait described in tomato, which is controlled by a single dominant gene known as the *Uniform ripening* (*U*) locus (MacArthur, 1934; Tanksley, 1992; Grandillo and Tanksley, 1996; Powell *et al*., 2012; Nguyen *et al*., 2014). Tomato green shoulder has been counter-selected by breeders, to improve fruit colour uniformity and facilitate maturity stage determination and harvesting (Powell *et al*., 2012). Since tomato and eggplant display many significant similarities in their domestication syndromes (Chapman, 2019), a similar selection mechanism may have occurred in eggplant since immature fruits contain higher levels of chlorophyll in the peel.

In this study, we identified *SmGLK2* as the candidate gene controlling eggplant FN by using different experimental and germplasm populations, combining wild and cultivated genepools: (i) identification of candidate genomic region in a multi-parent advanced generation inter-cross (MAGIC) population; (ii) validation through bulked segregant analysis by sequencing (BSA-Seq) in a contrasting F_2_ population; (iii) narrowing down the genomic region by fine-mapping in two <u>advanced</u> backcross (AB) sets; (iv) search for causative allelic variants in the candidate gene sequence by the resequencing of accessions contrasting for the presence (FN) or absence (fn) of netting in the fruit peel; (v) tracing the changes that the gene has undergone through the domestication process in a set of accessions from a diverse germplasm collection; (vi) evaluation of the effect of the main mutation identified at the protein level by Western blotting analysis; and (vii) observation of chlorophyll autofluorescence emission from different tissues by confocal microscopy. The identification of *SmGLK2* gene as responsible for the eggplant FN trait is useful to understand the genetic basis of fruit colouration in eggplant as well as for eggplant breeding.

## Materials and methods

### Plant materials

#### Multi-parent advanced generation inter-cross (MAGIC) population

A total of 420 individuals from the eggplant S3MEGGIC population developed by Mangino *et al*. (2022) were used to identify candidate genomic regions associated with the FN trait. This population was previously used for identifying *SmAPRR2* as the causative gene for uniform chlorophyll pigmentation in the eggplant fruit peel (Arrones *et al*., 2022). The S3MEGGIC population was obtained by inter-crossing seven *Solanum melongena* accession and one wild *S. incanum* INC accession (Figure 1A). Only the wild INC founder displayed the FN trait, while none of the seven *S. melongena* displayed this trait.

**Figure 1.**
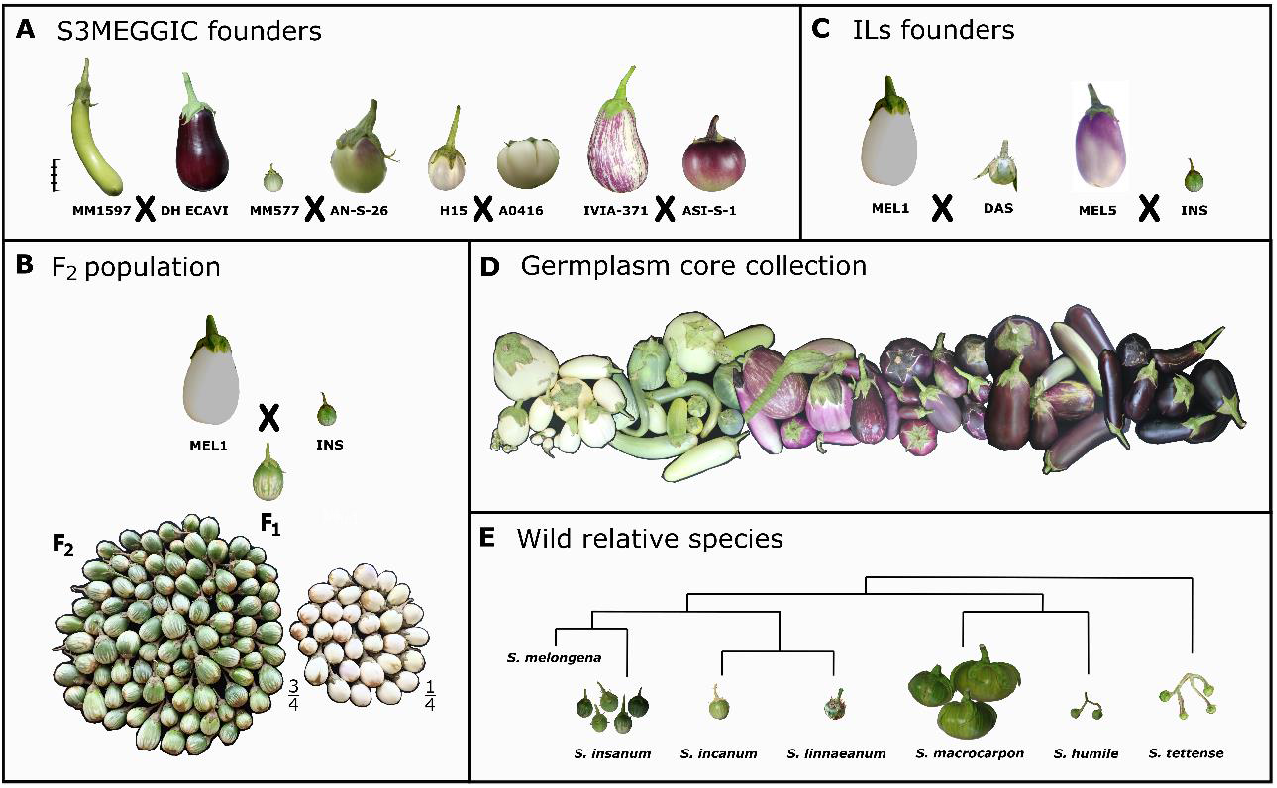
Plant materials used in the study. (A) The eight founders of the S3MEGGIC population developed by Mangino *et al*. (2022) that were inter-crosses in pairs to obtain the following generations. The scale bar represents 5 cm. (B) Parents, F_1_ and F_2_ segregating population for the fruit green netting (FN) trait. (C) Parents for the two interspecific introgression lines (ILs) populations: *Solanum melongena* (MEL1 and MEL5), *S. dasyphillum* (DAS), and *S. insanum* (INS). (D) The variability among the selected accessions from the eggplant G2P-SOL germplasm core collection. (E) Wild relative species of the common *S. melongena* represented in a phylogenetic tree, including *S. incanum, S. insanum, S. linneanum, S. macrocarpon, S. humile*, and *S. tettense*.

#### F_2_ population

The white-fruited accession *S. melongena* MEL1 and the green netting-fruited *S. insanum* INS were used as parents to develop an F_2_ population of 120 individuals segregating for the FN trait (Figure 1B). This population was used for the validation of the candidate genomic region. These parents were selected for the easier visualisation of the FN in the absence of anthocyanins (Figure S1).

#### Advanced Backcross individuals and selfings

Advanced backcross (AB) individuals from two programs of development of interspecific introgression lines (ILs) populations were used to narrow down the candidate genomic region responsible for the FN trait. One ILs population was developed by crossing one white *S. melongena* accession MEL1 and one accession of the wild *S. dasyphyllum* DAS, while the other ILs population was developed from the interspecific cross between one *S. melongena* MEL5, which presents fruit anthocyanin (Mangino *et al*., 2020), and the wild *S. insanum* INS (Plazas *et al*., 2020; Figure 1C). The two wild parents (DAS and INS) presented FN phenotype while none of the *S. melongena* (MEL1 and MEL5) had chlorophyll pigmentation in the fruit. For this study, AB individuals from the advanced stages of the ILs population development displaying the FN phenotype were selected. These AB individuals were used to confirm the FN candidate genomic region by locating the introgressed wild fragment by the single primer enrichment technology (SPET) high-throughput genotyping platform (Barchi *et al*., 2019a). The AB individuals with the shortest introgressed fragment were selfed to find recombinants in their progeny and further narrow down the candidate region.

#### Germplasm core collection and wild relative species

Genomic sequences of 178 *S. melongena* accessions (76 FN and 102 fn) available from the eggplant germplasm core collection established in the framework of the G2P-SOL project (http://www.g2p-sol.eu/G2P-SOL-gateway.html), were interrogated for the proposed candidate gene to find causative variants of the fn phenotype (Figure 1D). This core collection includes accessions used for developing the first eggplant pan-genome (Barchi *et al*., 2021). In addition, 22 wild relative species, including *S. insanum, S. incanum, S. linneanum, S. macrocarpon, S. humile*, and *S. tettense*, all with FN phenotype, were selected to confirm the putative candidate gene structure (Figure 1E).

### Cultivation Conditions

Seeds from the studied individuals were germinated in Petri dishes, following the protocol developed by Ranil *et al*. (2015). They were subsequently transferred to seedling trays in a climatic chamber under photoperiod and temperature regime of 16 h light (25 °C) and 8 h dark (18 °C). After acclimatization, plantlets were grown either in a pollinator-free benched glasshouse or an open field plot at the UPV campus in Valencia, Spain (GPS coordinates: latitude, 39° 28′ 55″ N; longitude, 0° 20′ 11″ W; 7 m above sea level). Plants were spaced 1.2 m between rows and 1.0 m within the row, fertirrigated using a drip irrigation system and trained with vertical strings (greenhouse) or bamboo canes (open field). Pruning was done manually to regulate vegetative growth and flowering. Phytosanitary treatments were performed when necessary. To accelerate fruit setting, inflorescences of plants grown in the greenhouse were vibrated with a mechanical vibrator.

### F_2_ population development for BSA-Seq analysis

The complete F_2_ population obtained by the interspecific cross between MEL1 and INS parents was phenotyped for the FN trait and a chi-square (χ^2^) test was performed to assess the goodness-of-fit to a 3:1 segregation model. This population was used for a bulked segregant analysis by sequencing (BSA-Seq). As the recommended size of each pool population is considered to be around 0.25 (Huang *et al*., 2022), 30 individuals were selected from each of the FN and fn pools.

Young leaves were harvested from both parents and the 60 F_2_ individuals selected by their different FN phenotypes. Genomic DNA was extracted using the silica matrix (SILEX) protocol (Vilanova *et al*., 2020) and checked for quality and integrity by agarose electrophoresis and NanoDrop (Thermo Fisher Scientific, Waltham, MA, USA) ratios (260/280 and 260/230), while its concentration was estimated with Qubit 2.0 Fluorometer (Thermo Fisher Scientific, Waltham, MA, USA). Pools were made by mixing an equal amount of DNA from each selected F_2_ individual (20 ng/µl) and shipped to Novogene (Novogene Europe, Cambridge, United Kingdom) where genomic libraries (PE 150, insert 350 bp) were constructed and sequenced.

All raw reads were trimmed using the fastq-mcf tool from the Ea-utils package (Aronesty, 2013) with “q 30 -l 50” parameters, and the overall quality was checked using FastQC v.0.12.1 (Andrews *et al*., 2010). Clean reads were mapped against the 67/3 v.3 eggplant reference genome (Barchi *et al*., 2019b) using the BWA-MEM v.0.7.17–r1188 with default parameters (Li and Durbin, 2009). The ΔSNP-index was estimated using the QTL-seq software v.2.2.3 (Takagi *et al*., 2013) from BAM files with “n1 30 -n2 30 -D 90 -d 5” options. The calculated ΔSNP-index were imported into R v.4.3.1 for the final plot.

### Fine-mapping of the AB introgressed fragments

ILs parents (MEL1, DAS, MEL5, and INS) and the selected AB individuals were high-throughput genotyped using the 5k eggplant SPET platform (Barchi *et al*., 2019a). The SNPs identified were filtered using the TASSEL software to retain the most reliable ones (minor allele frequency > 0.01, missing data < 20% and maximum marker heterozygosity < 80%) and the graphical visualization of the genotypes was performed by using Flapjack software (Milne *et al*., 2010).

To further narrow down the candidate region, AB individuals with the shortest wild introgressed fragment were selfed and the progeny was genotyped by High Resolution Melting (HRM) on a LightCycler 480 Real-Time PCR (Roche Diagnostics, Meylan, France) to identify recombinants with shorter introgressions. To design primers pairs spanning the region and detect recombination breakpoints with higher precision, the genome of the four ILs parents was resequenced (PE150, insert 300 bp, 24 Gb) at the Beijing Genomics Institute (BGI Genomics, Hong Kong, China). Trimming, quality check, mapping and SNP calling were performed against the 67/3 v.3 eggplant reference genome (Barchi *et al*., 2019b) as in Gramazio *et al*. (2019). Integrative Genomics Viewer (IGV) tool v.2.15.2 was used for the visual exploration and detection of variants among parental genome sequences (Robinson *et al*., 2023). Primers were designed to detect differential indel variants, which are listed in Table S1. The informative recombinants were grown in a pollinator-free benched glasshouse and phenotyped for FN.

### Candidate genes for FN and allelic variants

The genomic region narrowed down by fine mapping was explored for candidate genes, retrieved from the functional annotation of the 67/3 v.3 reference genome (Barchi *et al*., 2019b). Simultaneously, the whole genome resequencing data of the eight founders of the S3MEGGIC populations (Gramazio *et al*., 2019) and the four parents of the two ILs populations were interrogated for “high” impact allelic variants predicted by SnpEff (Cingolani *et al*., 2012) that were contrasting between FN and fn. The nucleotide sequence for the best candidate gene, the eggplant Golden 2-like MYB (*SmGLK2*), was retrieved by a BLASTx search (e-value cut-off of 1e-5) against the HQ-1315 and GUIQIE-1 eggplant reference genomes (Wei *et al*., 2020; Li *et al*., 2021). Additionally, transcriptomic data from *S. melongena* 67/3 (SRR3884608), *S. incanum* INC (SRR2289250), and *S. insanum* MM0686 accession (SRR8736646) were downloaded from the NCBI SRA database. Transcripts were mapped against the *SmGLK2* gene sequence of the 67/3 v.3 reference genome using RNA STAR (Dobin and Gingeras, 2015) and checked using the IGV tool. Gene sequence of the tomato ortholog (*SlGLK2*, Solyc10g008160.3) was also analysed to elucidate the *SmGLK2* gene structure. A conservative domain analysis was performed by assessing the NCBI conserved domain server (https://www.ncbi.nlm.nih.gov/Structure/cdd/wrpsb.cgi).

Allelic variants of the *SmGLK2* gene were also interrogated in whole-genome resequencing data of 178 *S. melongena* accessions from the eggplant G2P-SOL germplasm core collection and 22 wild relative species (Gramazio *et al*. 2019, Barchi et al 2021 and unpublished data) for which FN phenotyping was available. For this purpose, raw reads were trimmed by SOAPnuke software (Chen *et al*., 2018) with filter parameters “-l 20 -q 0.5 -n 0.03 -A 0.28” and aligned against de 67/3 v.3 eggplant reference genome (Barchi *et al*., 2019b) using the BWA-MEM program v.0.7.17–r1188 (Li and Durbin, 2009). Subsequently, the clean reads mapped in the genomic region of *SmGLK2* were extracted with samtools (Danecek *et al*., 2021) and *de novo* assembled using Megahit v.1.2.9 with default parameters (Li *et al*., 2015). The raw data of the assembled *SmGLK2* of the G2P-SOL collection is available at NCBI SRA (BioProject IDs PRJNA649091, PRJNA392603 and PRJNA977872). A multiple sequence alignment of the assembled sequences was then performed using the MAFFT program v.7 (Katoh and Hiroyuki, 2008) and the results were visualized in the Jalview alignment editor v.2.11.2.6 (Waterhouse *et al*., 2009). The resulting alignments were used to identify the candidate variants in the *SmGLK2* gene for the G2P-SOL core collection and wild accessions.

### Protein extraction and immunoblot analysis

Fruit peel tissues from eggplant accessions INS and MEL1 were harvested and ground with liquid nitrogen. Total proteins were extracted with extraction buffer [Tris-HCl 50 mM (pH 8), MgCl_2_ 10 mM, glicerol 5%, Triton X-100 0.1%, β-mercaptoethanol 14mM]. Samples were centrifuged twice at 13,000 rpm for 15 min to remove cell debris and recover the cleanest supernatant containing proteins. Fourteen mg of protein were denatured at 95 ºC for 5 min in reducing loading buffer and separated by 12% sodium dodecyl sulfatepolyacrylamide gel electrophoresis (SDS-PAGE; Sambrook and Russell, 2006). Proteins were transferred (30V 1h 4 ºC) to polyvinylidene fluoride membranes (Immobilon®-E PVDF Membrane) and blocked overnight at 4 ºC with 5% non-fat dry milk (Sveltesse®, Nestlé) and gentle shacking. To detect the gene product of *SmGLK2*, membranes were incubated with an anti-sera peptide based polyclonal rabbit GLK2-antibody (1:3,000) and a peroxidase-conjugated anti-rabbit antibody (1:10,000) (Merck, Darmstadt, Germany) for 1 h at room temperature in TBST [Tris-HCl 20 mM (pH 7.6), NaCl 20 mM, Tween20 0.1% v/v] supplemented with 2% non-fat dry milk. The anti-sera peptide based polyclonal rabbit GLK2-antibodies were raised against the *SmGLK2* N-terminal sequence (synthetic peptide VSPPLSYTNENENY, 5–18 aa; and NMKSKSKEAKKSSG, 70–83 aa) (Davids Biotechnologie GmbH, Regensburg, Germany). Peroxidase activity was developed in ECL Plus Western blotting detection reagents (Amersham, Buckinghamshire, England) and the chemiluminescence signal was captured with the LAS-1000 imaging system (Fujifilm, Tokio, Japan).

### Confocal microscopy

To determine differences in chloroplast number and structure between the netting and the uniform distribution of chlorophylls in the fruit peel as a result of *SmGLK2* and *SmAPRR2* expression, respectively, FN and fn tissues were analysed. Exploiting chlorophyll autofluorescence and emission spectra, confocal microscopy was used for monitoring chloroplast presence. Following the Confocal Laser Scanning Microscopy (CSLM) protocol developed by D’Andrea *et al*. (2014), thin sections of the pericarp were cut with a vegetable peeler or a single-edge razor blade. Samples were mounted on a microscope slide with a drop of water with the skin facing the coverslip. Samples were imaged in an AxioObserver 780 (Zeiss) confocal microscope using a 488 nm argon laser and a water-immersion 40x objective, and chlorophyll emission was recorded between 634 and 723 nm. For each sample, a stack of 10 nm was taken, taking one optical section every 1 µm, as well as a lambda scan between 500 and 700 nm with a bandwidth of 8 nm and a step size of 5 nm to verify that no other pigments were emitting in this range.

## Results

### BSA-seq for the FN trait

Through a GWAS in the S3MEGGIC population (Mangino *et al*., 2022), Arrones *et al*. (2022) identified a significant association peak for chlorophyll content in the eggplant fruit peel on chromosome 4 in the genomic region between 3.23 and 6.35 Mb (Figure 2A). However, no clear candidate gene was identified since the SnpEff software did not predict high-effect variants contrasting for the S3MEGGIC founders with and without chlorophylls in the fruit peel. The phenotyping of this population for fruit chlorophyll pigmentation was performed using a binary classification (presence/absence) without discriminating between uniform distribution or netting. This fact, together with previous studies relating this genomic region with the FN in eggplant (Tigchelaar, 1968; Doganlar *et al*., 2002; Frary *et al*., 2014), led us to hypothesize a possible relationship of this peak with the FN trait.

**Figure 2.**
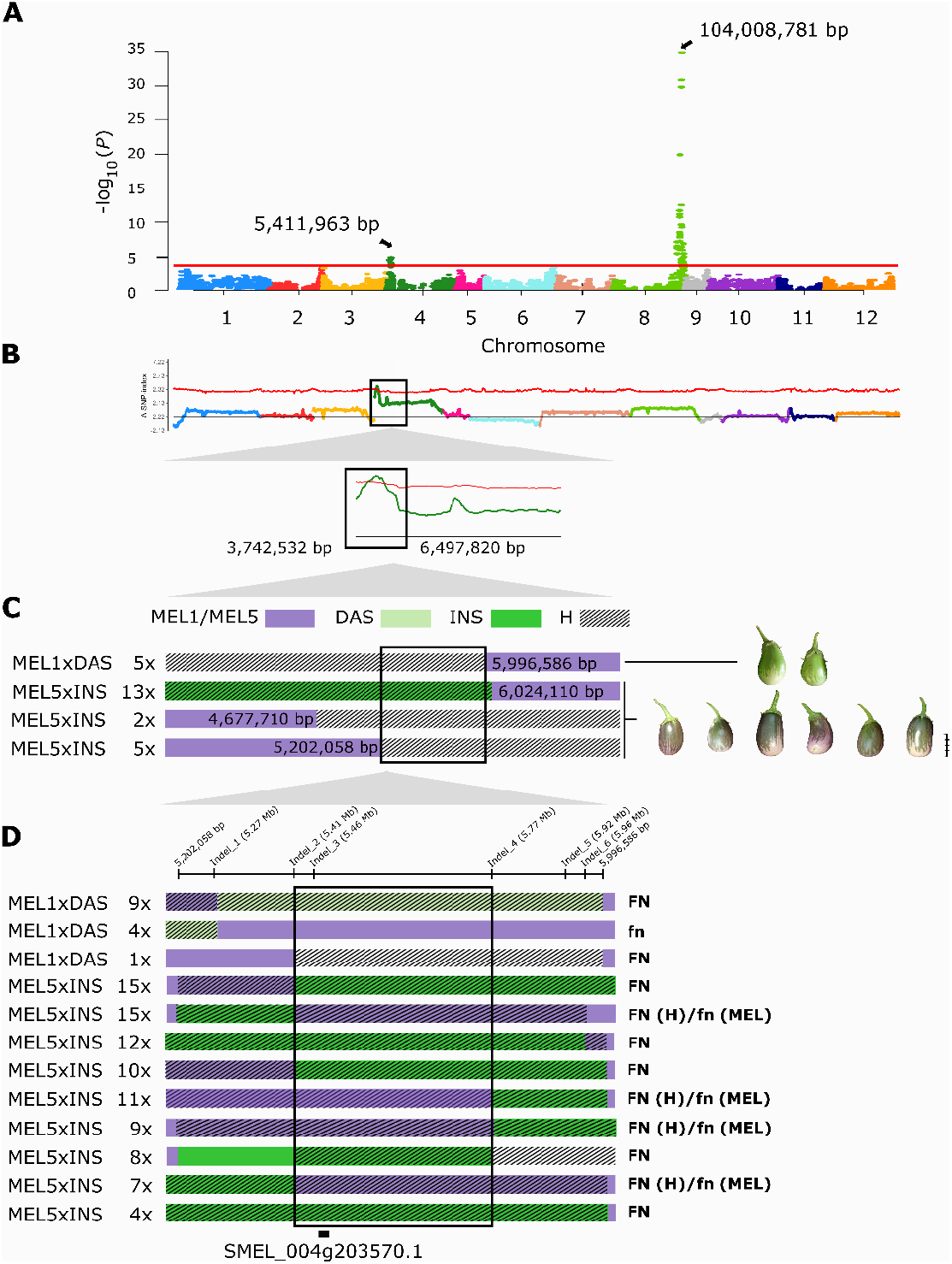
(A) Manhattan plot for fruit chlorophyll from the GWAS analysis in the S3MEGGIC population (Arrones *et al*., 2022). Arrows indicate the genome position of the significant highest peaks. The horizontal red line represents the FDR significance threshold at p = 0.05. (B) BSA-Seq results from the green netting-fruited and non-netting-fruited pools. The association significance threshold (ΔSNP-index) is represented with a red line. The black square delimits the significant candidate region. (C) Representation of the 25 selected advanced backcross (AB) individuals from two IL populations, MEL1 x DAS (5 accessions) and MEL5 x INS (20 accessions), covering the candidate region on chromosome 4. In purple the two *S. melongena* parents (MEL1 and MEL5) background, in green the introgressions coming from *S. dasyphyllum* (DAS) or *S. insanum* (INS), and with stripes the heterozygotes (H). On the right, the phenotype of the fruits obtained from the selected AB at a scale based on the real fruit size, all of them presenting the fruit green netting (FN). Scale bar represents 5 cm. (D) Representation of the 105 recombinant individuals coming from the 1,761 individuals generated by selfing of the AB individuals with the introgressed fragment in heterozygosis. The genomic indel positions used for the edges-to-core fine-mapping are specified on the top. The phenotype of the different individuals is indicated on the right. The position of the *SmGLK2* gene (SMEL_004g203570.1) is shown.

To confirm the candidate genomic region for the FN, an F_2_ population was developed by crossing MEL1 (fn) and INS (FN) parents. The FN trait displayed classical patterns of Mendelian inheritance of a single dominant gene in the F_1_ and F_2_ populations. Among the 120 F_2_ individuals, 74.17% (89 out of 120) were FN while 25.83% (31 out of 120) were fn (Figure 1C). Chi-square (χ^2^) analysis was in agreement with the 3:1 segregation with a value of 0.044 (d.f. = 1.0, P < 0.05). BSA-Seq was performed using 30 individuals from each of the FN and fn pools from the F_2_ segregating population. The DNA pools sequencing resulted in an average of 117 million 150 bp raw reads yielded per bulk with a mean coverage of 33X. About 9,436,870 high-quality SNPs differentiating between parental lines were identified. Of these, 2,192,129 SNPs were selected by QTLseq software to compute the ΔSNP-index with a 99% confidence interval identifying a unique significant region of 2.76 Mb on chromosome 4, between 3,742,532 bp and 6,497,820 bp, which was consistent with the results obtained in the S3MEGGIC population (Figure 2B, Table S2).

### Fine-mapping of the candidate genomic region

Since two AB populations segregating for the FN trait were available, both were used for further inspecting the candidate genomic region identified by GWAS and BSA-seq analyses. Specifically, five AB individuals from the MEL1 × DAS set and 20 AB individuals from the MEL5 × INS were selected, all showing the FN phenotype. All of the 25 AB individuals presented a wild DAS or INS introgression in a *S. melongena* background (MEL1 and MEL5, respectively) at the beginning of chromosome 4. The size and the physical location of the wild introgressed fragments varied according to the recombination and the selection made in the previous generation. The genotyping of these AB individuals by the eggplant SPET platform allowed to narrow down the candidate region to 0.80 Mb, between 5,202,058 bp and 5,996,586 bp (Figure 2C). There were approximately 48 putative candidate genes under this narrowed region (Table S3).

To further delimit this region, 1,761 individuals were generated by selfing the AB individuals with the shortest wild introgressed fragment and screened following the edges-to-core approach. As a result, 105 recombinant individuals were identified (Table S4) and based on their genotype and phenotype, the region of interest was reduced to 0.36 Mb, between 5,412,659 bp and 5,774,878 bp of chromosome 4 (Figure 2D). The number of putative candidate genes under this region was reduced to 14 genes (highlighted in Table S3).

### Identification of a responsible candidate gene

The 14 candidate genes annotated according to the 67/3 v.3 reference genome (Barchi *et al*., 2019b) were interrogated for “high” impact allelic variants following SnpEff analysis, but none of the genes carrying such variants were consistent with the contrasting phenotypes. However, we decided to further interrogate the Golden 2-like MYB gene (*SmGLK2*, SMEL_004g203570.1, 5,457,658 – 5,461,306 bp) which was within the confidence interval, since it had previously been described as a positive regulator of chloroplast development and pigment accumulation in other Solanaceae species (Powell *et al*., 2012; Brand et al, 2014; Nguyen *et al*., 2014).

Although no high-effect variants were predicted by SnpEff in the coding sequence of *SmGLK2*, some intronic variants were shared by the FN S3MEGGIC founders and ILs parents (INC, DAS, and INS), compared to fn ones. Studying the gene structure in the other available eggplant reference genomes HQ-1315 and GUIQIE-1 (Wei *et al*., 2020; Li *et al*., 2021), it was observed that the *SmGLK2* gene model was split into two different genes (Smechr0400347-8 and EGP26827-8, respectively) between the second and third exon (Figure 3A). Furthermore, slight differences between *SmGLK2* gene structure compared to its tomato ortholog were identified (*SlGLK2*, Solyc10g008160.3). The *SlGLK2* 5’-UTR was divided into three different exons, the latter two coinciding with the two first exons of the *SmGLK2*. In addition, one extra exon was observed between the eggplant second and third exons (Figure 3A). Then, different transcriptomic data were analysed to reveal whether a gene mis-annotation existed in the eggplant reference genomes. When analysing the 67/3, INC, and MM0686 transcriptomes against the *SmGLK2* gene sequence of the 67/3 v.3 reference genome (Barchi *et al*., 2019b), some transcripts also aligned with the intronic regions between the second and third exon (Figure 3B), suggesting an incorrect annotation in the available eggplant reference genomes, all of them developed from fn accessions. We therefore propose the presence of an extra exon between the second and third exon for the 67/3 v.3 reference genome and the merging of the two annotated genes in the HQ-1315 and GUIQIE-1 eggplant reference genomes into a single one (Wei *et al*., 2020; Li *et al*., 2021; Figure 3C).

**Figure 3.**
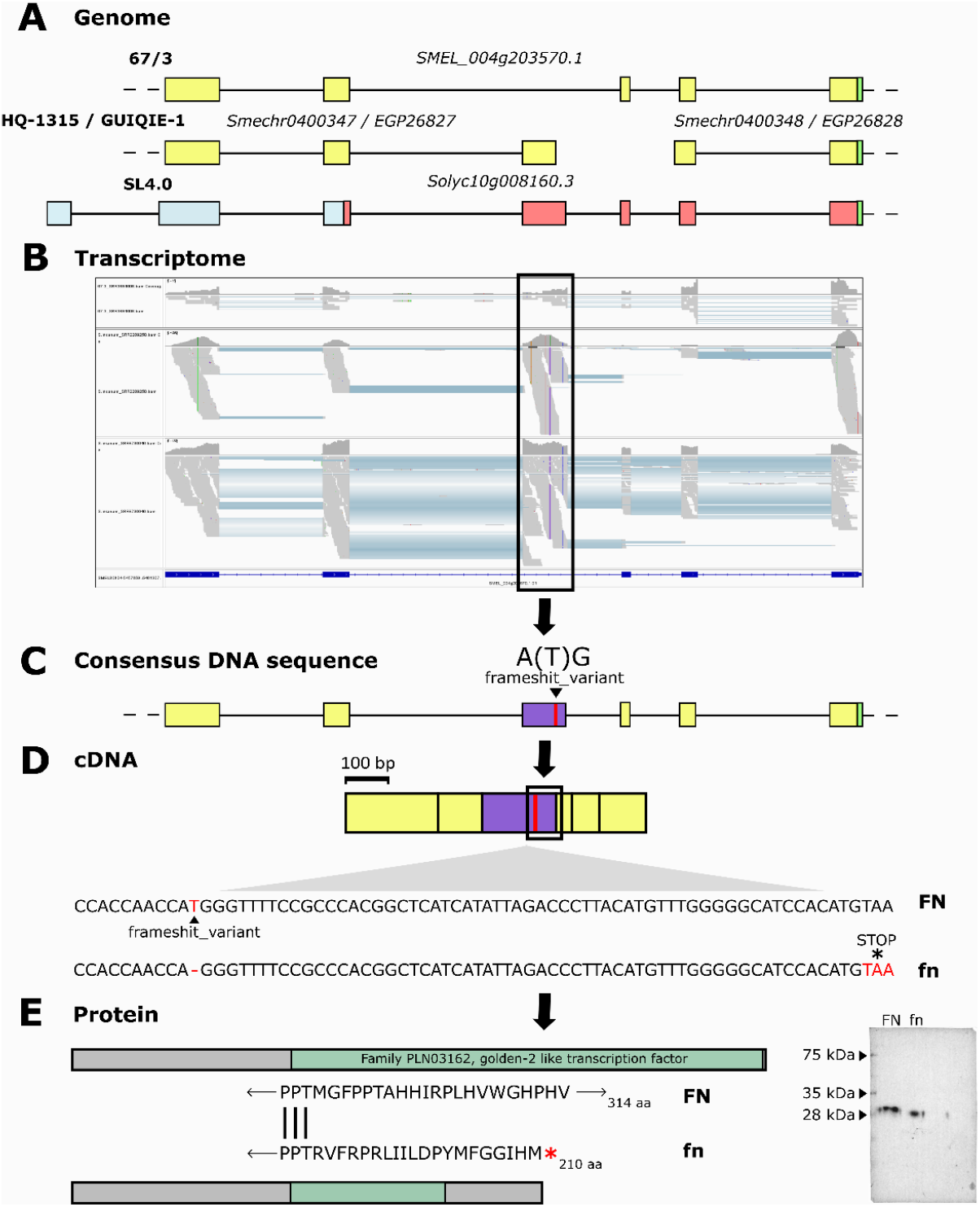
From the genomic sequence of the *SmGLK2* gene to protein structure. (A) *SmGLK2* annotated gene structure in 67/3, HQ-1315, and GUIQIE-1 eggplant reference genomes and SL4.0 tomato genome (5’-UTR in blue, eggplant annotated exons in yellow, tomato annotated exons in red, and 3’-UTR in green). (B) The *S. melongena* 67/3 (SRR3884608), *S. incanum* INC (SRR2289250), and *S. insanum* MM0686 accession (SRR8736646) transcripts aligned against the *SmGLK2* gene sequence from the 67/3 v.3 eggplant reference genome and visualized in the IGV tool (Robinson *et al*., 2023). The candidate structural variation identified is indicated with a purple line. (C) The suggested *SmGLK2* gene structural annotation as a consensus of previous information with the extra proposed exon in purple. The identified structural variation in the gene sequence is indicated with a black arrowhead and a red line. (D) Comparison of mRNA sequences for presence (FN) or absence (fn) of the fruit green netting trait. The premature stop codon downstream of the indel is indicated with an asterisk. (E) Comparison of the protein structure and sequence around the indel site for FN and fn. The golden-2 like transcription factor domain is indicated in green. On the right, a Western blot showing the difference in apparent molecular mass of the GLK2 protein from FN and fn tissues.

### Identification of causative allelic variants

In the suggested additional exon, a frameshift mutation was identified when comparing FN accessions against the 67/3 v.3 eggplant reference genome (Barchi *et al*., 2019b), which was derived from an fn accession. This mutation consisted of a small insertion A(T)G in the 5,459,673 bp position (Figure 3C). In the presence of this insertion, a full-length GLK2 protein is generated, while in its absence, a premature stop codon leads to a truncated GLK2 protein (Figure 3D and 3E). Analysing the resequencing data, all FN parents (the wild INC, DAS, and INS) presented the T insertion, while the fn parents were identical to the 67/3 reference. The absence or presence of this insertion was also confirmed in the 67/3 (fn), INC (FN), and MM0686 (FN) transcripts.

To further study the *SmGLK2* allelic variants responsible for the loss of the FN trait over the course of domestication, a diverse germplasm collection was analysed. The aim was to trace the changes that the gene had undergone from wild to cultivated species. Therefore, we selected 178 *S. melongena* accessions from the eggplant G2P-SOL germplasm core collection, 76 FN and 102 fn, and 22 wild relative species, all of which showed the FN phenotype (Table S5). All 76 FN accessions from the G2P-SOL core collection presented the T insertion. However, only 97 out of 102 fn accessions showed the absence of the T insertion, similar to 67/3. The remaining five accessions exhibited a different haplotype, with two new frameshift mutations: a 68 bp deletion on exon 1 from 5,457,779 – 5,457,847 bp and a 41 bp deletion on exon 5 from 5,461,264 – 5,461,305 bp. The 22 wild relative species analysed also presented the T insertion, indicating a complete and fully functional *GLK2* gene.

### Differences in GLK2 protein structure

Analysis of the full-length amino acid sequence of *SmGLK2* in the NCBI conserved domain server, confirmed that the premature stop codon generated by the absence of the T insertion was translated into a truncated protein in fn accessions. Precisely, 148 aa of the golden-2-like transcription factor domain at the C-terminal region were lost. Western blot analysis with a polyclonal rabbit GLK2 antibody showed a differential molecular mass banding pattern of the target protein for INS (FN) and MEL1 (fn) samples. A difference in band migration was observed, with the apparent molecular masses of full-length and truncated versions of GLK2 being of approximately 30.5 kDa for FN and 28.75 kDa for fn, respectively (Figure 3E).

### Cytological observation

Confocal microscopy was used to determine differences in chloroplast number and structure in FN and fn fruits. Fruit peel from areas displaying netting (FN) and uniform green coloration (fn) exhibited important differences (Figure 4). In the FN sample coming from the proximal part of a fruit with dark green netting, chloroplasts showed higher chlorophyll content. Instead, in the fn sample chloroplasts showed low levels of chlorophylls. Overall, our results indicate that the presence of a full-length GLK2 protein leads to the accumulation of higher amounts of chlorophylls, which results in the dark-green netted pattern in eggplant fruits.

**Figure 4.**
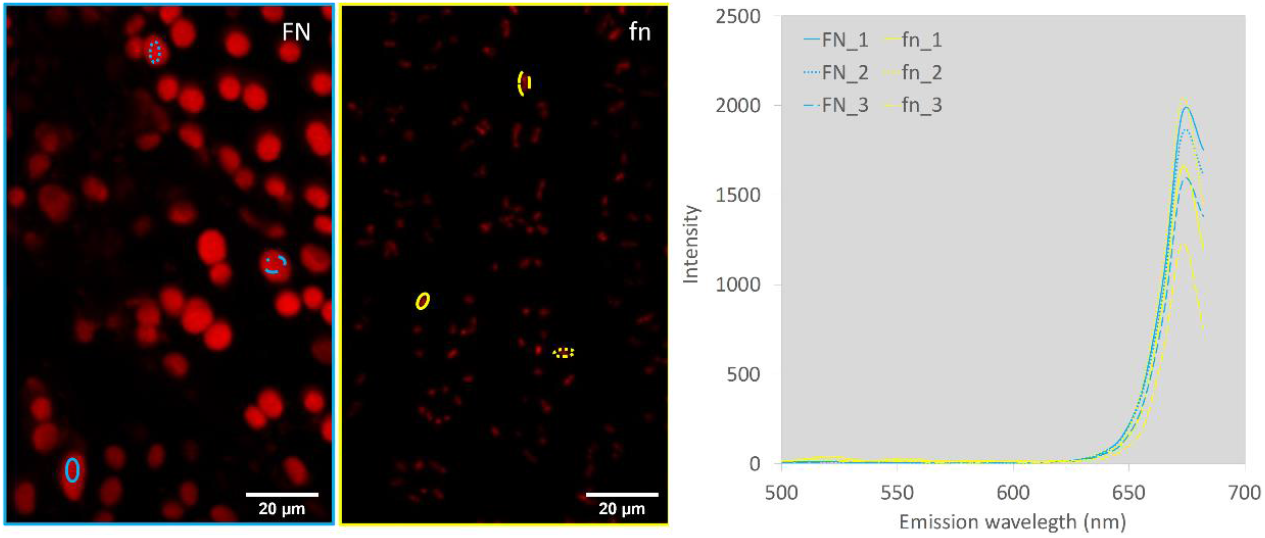
Confocal images of pericarp samples showing presence (FN) or absence (fn) of the fruit green netting trait. The FN sample comes from the proximal part of a fruit showing a dark green netting (*SmGLK2* expression), while the fn sample comes from a fruit with uniform distribution of chlorophylls (*SmAPRR2* expression). On the right, fluorescence emission spectra from FN and fn samples after excitation at 488 nm. Fluorescence intensity is represented relative to the total fluorescence of the sample.

## Discussion

Eggplant domestication resulted in a diversification of fruit colour at the physiologically unripe stage, characteristic of commercial maturity. However, in recent times, modern breeding has resulted in a predominance of purple and dark purple colours, especially in Western markets (Taher *et al*., 2017; Page *et al*., 2019; Page *et al*., 2021). Consumers demand new vegetable products with different and unusual aesthetic characteristics, broadening the consumers’ options and fostering vegetable consumption (Di Gioia *et al*., 2020). In this study, we have successfully harnessed different plant materials, combining germplasm and classical bi-parental (F_2_) populations with advanced bi- (ILs) and multi-parent (MAGIC) populations, for the identification of the genes underlying FN in eggplant. This is interesting as it increases the diversity of the eggplant fruit colour palette.

Research on the genetics of FN in eggplants, also known as green reticulation or variegation, has been limited. Daunay *et al*. (2004) associated FN with a trait under monogenic dominant control following an F_2_ proportion of 3:1. We confirmed the dominant hypothesis by the development of the F_1_ hybrid and F_2_ population from an FN × fn cross. In the published studies on this trait, *S. linneanum* was used as a parental donor since FN is commonly present in this wild species (Doganlar *et al*., 2002; Daunay *et al*., 2004; Frary *et al*., 2014). Although a few commercial varieties display the FN trait, it has been counter-selected during domestication because the green colour was erroneously related to immaturity (Page *et al*., 2019). As occurs with other domestication traits, fn is recessive to the dominant allele of the wild species (Doganlar *et al*., 2002; Frary *et al*., 2014). This supports the paradigm that domestication usually involves the loss of gene function or regulation (Lester and Hasan, 1991).

In our previous study, through a GWAS in the S3MEGGIC population, we identified significant associations for the presence of chlorophyll pigmentation in the eggplant fruit peel (Arrones *et al*., 2022). The Manhattan plot revealed one major peak on chromosome 8, which resulted in the identification of a gene identified as similar to ARABIDOPSIS PSEUDO RESPONSE REGULATOR2 (*APRR2*, SMEL_008g315370.1). The *SmAPRR2* gene was suggested as the best candidate gene for uniform fruit chlorophyll pigmentation. The GWAS also revealed a minor peak, although significant, at the beginning of chromosome 4. This region was rejected as SnpEff did not predict contrasting high-effect variants among the S3MEGGIC founders. Since previous studies identified a QTL related to FN on chromosome 4 explaining 67-78% of the variation (Doganlar *et al*., 2002; Daunay *et al*., 2004; Frary *et al*., 2014), we decided to further investigate this genomic region. Moreover, the minor peak could have been obtained by phenotyping imprecisions as no discrimination between netting or uniform distribution of chlorophylls in the Arrones *et al*. (2022) study. These results together with a BSA-Seq of an F_2_ population segregating for the trait and the fine mapping of two ABs populations, allowed us to confirm the candidate genomic region for FN and to narrow it down to 0.36 Mb. None of the 14 genes under this region presented variants consistent with the FN and fn phenotypes. However, one of these genes annotated as similar to Golden 2-like MYB (*SmGLK2*, SMEL_004g203570.1) was studied as the best candidate. This gene belongs to the widely conserved GARP family of MYB transcription factors whose functions are reported to be involved in chloroplast development in different species such as *Physcomitrella patens* (moss), *Zea mays* (maize), and *Arabidopsis thaliana* (Hall *et al*., 1998; Rossini *et al*., 2001; Fitter *et al*., 2002; Yasumura *et al*., 2005).

Through different complementary analyses, we identified an erroneous annotation of the *SmGLK2* in the available reference genomes, all of them developed from accessions with fn phenotype (Barchi *et al*., 2019b; Wei *et al*., 2020; Li *et al*., 2021). An improved annotation of the *SmGLK2* gene was achieved thanks to the availability of resequencing and transcriptomic data. As a result, we found an extra coding exon where we identified a small insertion. The complete sequence of the *SmGLK2* gene is given by the presence of this insertion, while its absence results in a frameshift mutation and a premature stop codon in the mRNA sequence. This was the reason for the fn phenotypes in our populations. This could also explain the HQ-1315 and GUIQIE-1 eggplant reference genomes annotation of *SmGLK2* as two different genes (Wei *et al*., 2020; Li *et al*., 2021). Furthermore, we verified the effects of this mutation at the protein level by Western blotting analysis. The presence of the small insertion resulted in a complete and fully functional *SmGLK2* gene product, while its absence resulted in a truncated golden-2-like transcription factor domain leading to a protein of smaller molecular weight.

Reverse genetics approaches, such as over-expression of the candidate genes or knock-out by CRISPR/Cas are the optimal gene validation techniques. However, due to the recalcitrance of eggplant to genetic transformation (García-Fortea *et al*., 2020; Mir *et al*., 2021), there is a need for alternative indirect validation methods in this crop until new, genotype-independent genetic transformation protocols become available. Validation through diverse experimental populations and the screening of large germplasm collections can provide a good alternative (Arrones *et al*., 2022). For this reason, a set of *S. melongena* accessions from the eggplant G2P-SOL germplasm core collection and wild species, including *S. insanum, S. incanum, S. linneanum, S. macrocarpon, S. humile*, and *S. tettense*, were analysed. This methodology also allows for tracing the evolutionary changes that the gene has undergone. The complete *SmGLK2* gene sequence was confirmed in the wild relative species, all of them with FN phenotype. The small insertion identified was validated in the G2P-SOL germplasm core collection, contrasting genotyping and phenotyping data. Besides, two additional, rare frameshift mutations responsible for the fn phenotype were identified. This indicates that the lack of the FN trait has arisen and been selected independently several times during eggplant domestication and diversification.

All these results together provide strong evidence of *SmGLK2* gene as the gene controlling the FN trait. These results are also supported by previous studies performed in other Solanaceae species. In pepper, *CaGLK2* has also been described as a major gene controlling chlorophyll content and chloroplast compartment size in immature fruit (Brand et al, 2014). Three null mutations resulting in the appearance of premature stop codons and leading to truncated proteins were identified as responsible for a reduction in the chlorophyll content of pepper fruits. In tomato, *SlGLK2* has been related to the green shoulder trait, also referred to as the *Uniform ripening* (*U*) locus, which is very similar to FN in eggplant (MacArthur, 1934; Tanksley, 1992; Grandillo and Tanksley, 1996; Powell *et al*., 2012). It has been demonstrated that the overexpression of *SlGLK2* increases chloroplast number and size, producing homogeneously dark-green fruits with enhanced nutritional quality (Powell *et al*., 2012; Nguyen *et al*., 2014). An A insertion causing a frameshift and a premature stop codon encoding for a truncated protein was identified as responsible of the *u* phenotype (Powell *et al*., 2012). Moreover, Nadakuduti *et al*. (2014) suggested that *SlGLK2* and *SlAPRR2* act independently as key transcription factors to directly activate genes involved in fruit chloroplast development in tomato. Likewise, the results obtained by confocal imaging of chlorophyll autofluorescence also support different roles for *SmGLK2* and *SmAPRR2* leading to two independent traits. Ultrastructural analysis of chloroplasts in netted areas of tomato fruits showed more developed chloroplasts, with more grana in the stroma than those of non-netted tomatoes (Powell *et al*., 2012; Nguyen *et al*., 2014). This is coherent with our observation of higher amounts of chlorophylls in eggplant netted areas. Overall, these results indicate that *SmGLK2* expression leads to more developed chloroplasts with higher chlorophyll accumulation. This may also be the reason why the green colour coming from the netting is persistent and remains throughout ripening, until turning yellow when over ripe, while chlorophylls from uniform pigmentation fade rapidly and give rise to yellow pigments (Page *et al*., 2021).

By using multiple *in silico* and *in vivo* methodologies we have found that *SmGLK2* gene is responsible for the FN trait, with a 100% association between disruptive mutations in this gene and fn phenotypes. The eggplant FN has important implications for fruit visual quality particularly because eggplant is commercialized when it is still physiologically immature, and the expression of the trait is very intense in the proximal part of the fruit. The rescue of this domestication trait for future eggplant breeding programmes could be interesting to widen the diversity of the fruit colour palette. In addition, due to a higher concentration of functional chloroplasts and chlorophyll accumulation, *SmGLK2* could also be a valuable target for enhancing fruit nutritional quality.

## Acknowledgements

This work has been funded by grants PID2021-128148OB-I00 funded by MCIN/AEI/10.13039/501100011033/ and by “ERDF A way of making Europe”, PDC2022-133513-I00 funded by MCIN/AEI/10.13039/501100011033/ and European Union Next Generation EU/ PRTR, CIPROM/2021/020 from Conselleria d’Innovació, Universitats, Ciència i Societat Digital (Generalitat Valenciana, Spain), and by European Union’s Horizon 2020 Research and Innovation Programme under Grant Agreement No. 677379 (G2P-SOL project: Linking genetic resources, genomes and phenotypes of Solanaceous crops). AA is grateful to Spanish Ministerio de Ciencia, Innovación y Universidades for a predoctoral (FPU18/01742) contract. VB-F is grateful to Generalitat Valenciana for grant INVEST/2022/146, funded by European Union, Next Generation EU. SM would like to thank funding support from UPV and the Spanish Ministerio de Universidades under the program María Zambrano funded by the European Union Next Generation-EU. PG is grateful to Spanish Ministerio de Ciencia e Innovación for a post-doctoral grant (RYC2021–031,999-I) funded by MCIN/AEI /10.13039/501,100,011,033 and the European Union through NextGenerationEU/PRTR.

## Contributions

JP, PG, and SV conceived the idea and supervised the study. AA, MP, and PG performed the field trials. AA, VB-F, PG, and SV performed the analysis of the S3MEGGIC, F_2_ and AB populations and the *GLK2* gene structure. EP, LB, and GG performed the analyses of the G2P-SOL core collection. AA and SM performed confocal and protein analysis. AA and PG prepared a first draft of the manuscript and the rest of the authors reviewed and edited the manuscript. All authors have read and agreed to the published version of the manuscript.

## Data availability statement

The data presented in the study are deposited in the NCBI SRA repository, BioProject IDs PRJNA649091, PRJNA392603 and PRJNA977872.

## Supplementary information

**Figure S1**. Appearance of eggplant fruits in presence (FN) or absence (fn) of the green netting and in combination with chlorophylls or/and anthocyanins.

**Table S1**. List of primers used in this study.

**Table S2**. Results obtained for the ΔSNP-index calculations on chromosome 4 using the QTL-seq software. Highlighted in yellow, the significant region between 3.74-6.50 Mb.

**Table S3**. List of putative candidate genes for the eggplant fruit green netting (FN) under 5.20-5.99 Mb region on chromosome 4. Genes are divided into blocks according to the primers designed for the candidate region fine-mapping. Highlighted in yellow, the 14 candidate genes under the 5.41-5.77 Mb delimited region.

**Table S4**. List of *Solanum melongena* accessions from the eggplant G2P-SOL germplasm core collection and wild relative species, indicating their phenotype for the fruit green netting presence (FN) or absence (fn), and their genotype for the identified allelic variants.

**Table S5**. Genotype of the 1,761 accessions used for the edges-to-core fine-mapping approach, indicating for each indel position if the individual was heterozygous (H) or homozygous for *S. melongena* (MEL1 or MEL5), *S. dasyphyllum* (DAS), or *S. insanum* (INS).

## References

Andrews S. FastQC: a quality control tool for high throughput sequence data. 2010.

Aronesty E. Comparison of sequencing utility programs. The open bioinformatics J 2013; 7: 1–8 DOI: 10.2174/1875036201307010001.

Arrones A, Mangino G, Alonso D et al. Mutations in the SmAPRR2 transcription factor suppressing chlorophyll pigmentation in the eggplant fruit peel are key drivers of a diversified colour palette. Front Plant Sci 2022; 13, DOI: 10.3389/fpls.2022.1025951.

Barchi L, Acquadro A, Alonso D et al. Single Primer Enrichment Technology (SPET) for high-throughput genotyping in tomato and eggplant germplasm. Front Plant Sci 2019a; 10: 1005, DOI: 10.3389/fpls.2019.01005.

Barchi L, Pietrella M, Venturini L et al. A chromosome-anchored eggplant genome sequence reveals key events in Solanaceae evolution. Sci Rep 2019b; 9: 11769, DOI: 10.1038/s41598-019-47985-w.

Barchi L, Rabanus-Wallace MT, Prohens J et al. Improved genome assembly and pan-genome provide key insights into eggplant domestication and breeding. Plant J 2021; 107: 579–96, DOI: 10.1111/tpj.15313.

Blanke MM, Lenz F. Fruit photosynthesis. Plant, Cell & Environment 1989; 12, 31–46, DOI:doi.org/10.1111/j.1365-3040.1989.tb01914.x.

Brand A, Borovsky Y, Hill T et al. CaGLK2 regulates natural variation of chlorophyll content and fruit color in pepper fruit. Theor Appl Genet 2014; 127: 2139–48, DOI: 10.1007/s00122-014-2367-y.

Chapman MA. Introduction: The importance of eggplant. The Eggplant Genome, Springer, Cham. 2019; 1–10, DOI: 10.1007/978-3-319-99208-2_1.

Chen Y, Chen Y, Shi C et al. SOAPnuke: A MapReduce acceleration-supported software for integrated quality control and preprocessing of high-throughput sequencing data. Gigascience 2018; 7: 1–6, DOI: 10.1093/gigascience/gix120.

Cingolani P, Platts A, Wang LL et al. A program for annotating and predicting the effects of single nucleotide polymorphisms, SnpEff. Fly (Austin) 2012; 6: 80–92, DOI: 10.4161/fly.19695.

D’Andrea L, Amenós M, Rodríguez-Concepción M. Confocal laser scanning microscopy detection of chlorophylls and carotenoids in chloroplasts and chromoplasts of tomato fruit. Plant Isoprenoids: Methods in Molecular Biology. Vol 1153. 2014, 227–32, DOI: 10.1007/978-1-4939-0606-2_16.

Danecek P, Bonfield JK, Liddle J et al. Twelve years of SAMtools and BCFtools. Gigascience 2021; 10, DOI: 10.1093/gigascience/giab008.

Daunay M-C, Aubert S, Frary A et al. Eggplant (Solanum melongena) fruit colour: pigments, measurements and genetics. XIIth Meeting on Genetics and Breeding of Capsicum and Eggplant. 2004, 108–16.

Di Gioia F, Tzortzakis N, Rouphael Y et al. Grown to be blue—antioxidant properties and health effects of colored vegetables. Part II: Leafy, fruit, and other vegetables. Antioxidants 2020; 9, DOI: 10.3390/antiox9020097.

Dobin A, Gingeras TR. Mapping RNA-seq reads with STAR. Curr Protoc Bioinforma 2015; 51: 11.14.1-11.14.19, DOI: 10.1002/0471250953.bi1114s51.

Doganlar S, Frary A, Daunay M-C et al. Conservation of gene function in the Solanaceae as revealed by comparative mapping of domestication traits in eggplant. Genetics 2002; 161: 1713–26, DOI: 10.1093/genetics/161.4.1713.

Fitter DW, Martin DJ, Copley MJ et al. GLK gene pairs regulate chloroplast development in diverse plant species. Plant J 2002; 31: 713–27, DOI: 10.1046/j.1365-313X.2002.01390.x.

Frary A, Frary A, Daunay MC et al. QTL hotspots in eggplant (Solanum melongena) detected with a high resolution map and CIM analysis. Euphytica 2014; 197: 211–28, DOI: 10.1007/s10681-013-1060-6.

Frary A, Doğanlar S. Comparative genetics of crop plant domestication and evolution. Turkish J Agric For 2003; 27: 59–69.

Fuller DQ. Contrasting patterns in crop domestication and domestication rates: Recent archaeobotanical insights from the old world. Ann Bot 2007; 100: 903–24, DOI: 10.1093/aob/mcm048.

García-Fortea E, Lluch-Ruiz A, Pineda-Chaza BJ et al. A highly efficient organogenesis protocol based on zeatin riboside for in vitro regeneration of eggplant. BMC Plant Biol 2020; 20, DOI: 10.1186/s12870-019-2215-y.

Gramazio P, Yan H, Hasing T et al. Whole-genome resequencing of seven eggplant (Solanum melongena) and one wild relative (S. incanum) accessions provides new insights and breeding tools for eggplant enhancement. Front Plant Sci 2019; 10, DOI: 10.3389/fpls.2019.01220.

Grandillo S, Tanksley SD. QTL analysis of horticultural traits differentiating the cultivated tomato from the closely related species Lycopersicon pimpinellifolium. Theor Appl Genet 1996; 92: 935–51, DOI: 10.1007/BF00224033.

Hall LN, Rossini L, Cribb L et al. GOLDEN 2: A novel transcriptional regulator of cellular differentiation in the maize leaf. Plant Cell 1998; 10: 925–36, DOI: 10.1105/tpc.10.6.925.

Huang L, Tang W, Wu W. Optimization of BSA-seq experiment for QTL mapping. G3 Genes, Genomes, Genet 2022; 12, DOI: 10.1093/G3JOURNAL/JKAB370.

Katoh K, Toh H. Recent developments in the MAFFT multiple sequence alignment program. Brief Bioinform 2008; 9: 286–98, DOI: 10.1093/bib/bbn013.

Lester RN, Hasan SMZ. Origin and domestication of the brinjal eggplant, Solanum melongena, from S. incanum, in Africa and Asia. Solanaceae III: taxonomy, chemistry, evolution. The Linnean Society of London, London, UK, 1991; 369–387.

Li D, Qian J, Li W et al. A high-quality genome assembly of the eggplant provides insights into the molecular basis of disease resistance and chlorogenic acid synthesis. Mol Ecol Resour 2021; 21: 1274–86, DOI: 10.1111/1755-0998.13321.

Li D, Liu CM, Luo R et al. MEGAHIT: An ultra-fast single-node solution for large and complex metagenomics assembly via succinct de Bruijn graph. Bioinformatics 2015; 31: 1674–6, DOI: 10.1093/bioinformatics/btv033.

Li H, Durbin R. Fast and accurate short read alignment with Burrows-Wheeler transform. Bioinformatics 2009; 25: 1754–60, DOI: 10.1093/bioinformatics/btp324.

MacArthur JW. Linkage groups in the tomato. J Genet 1934; 29: 123–33, DOI: 10.1007/BF02981789.

Mangino G, Arrones A, Plazas M et al. Newly developed MAGIC population allows identification of strong associations and candidate genes for anthocyanin pigmentation in eggplant. Front Plant Sci 2022; 13, DOI: 10.3389/fpls.2022.847789.

Mangino G, Plazas M, Vilanova S et al. Performance of a set of eggplant (Solanum melongena) lines with introgressions from its wild relative S. incanum under open field and screenhouse conditions and detection of QTLs. Agronomy 2020; 10, DOI: 10.3390/agronomy10040467.

Milne I, Shaw P, Stephen G et al. Flapjack-graphical genotype visualization. Bioinformatics 2010; 26: 3133–4, DOI: 10.1093/bioinformatics/btq580.

Mir R, Calabuig-Serna A, Seguí-Simarro JM. Doubled haploids in eggplant. Biology (Basel) 2021; 10, DOI: 10.3390/biology10070685.

Nadakuduti SS, Holdsworth WL, Klein CL et al. KNOX genes influence a gradient of fruit chloroplast development through regulation of GOLDEN2-LIKE expression in tomato. Plant J 2014; 78: 1022–33, DOI: 10.1111/tpj.12529.

Nguyen C V., Vrebalov JT, Gapper NE et al. Tomato GOLDEN2-LIKE transcription factors reveal molecular gradients that function during fruit development and ripening. Plant Cell 2014; 26: 585–601, DOI: 10.1105/tpc.113.118794.

Page AML, Daunay M-CR, Aubriot X et al. Domestication of eggplants: A phenotypic and genomic insight. The Eggplant Genome. 2019, 193–212, DOI: 10.1007/978-3-319-99208-2_12.

Page AML, Chapman MA. Identifying genomic regions targeted during eggplant domestication using transcriptome data. J Hered 2021; 112: 519–25, DOI: 10.1093/jhered/esab035.

Plazas M, Gramazio P, Vilanova S et al. Introgression breeding from crop wild relatives in eggplant landraces for adaptation to climate change. Crop Wild Relat 2020, 32.

Powell ALT, Nguyen C V., Hill T et al. Uniform rippening encodes a golden 2-like transcription factor regulating tomato fruit chloroplast development. Science 2012; 336: 1708–11, DOI: 10.1126/science.1221863.

Ranil RHG, Niran HML, Plazas M et al. Improving seed germination of the eggplant rootstock Solanum torvum by testing multiple factors using an orthogonal array design. Sci Hortic (Amsterdam) 2015; 193: 174–81, DOI: 10.1016/j.scienta.2015.07.030.

Robinson JT, Thorvaldsdóttir H, Turner D et al. igv.js: an embeddable JavaScript implementation of the Integrative Genomics Viewer (IGV). Bioinformatics 2023; 39: btac830, DOI: 10.1101/2020.05.03.075499.

Rossini L, Cribb L, Martin DJ et al. The maize Golden2 gene defines a novel class of transcriptional regulators in plants. Plant Cell 2001; 13: 1231–44, DOI: 10.1105/tpc.13.5.1231.

Sambrook J, Russell DW. SDS-polyacrylamide gel electrophoresis of proteins. Cold Spring Harbor Protoc 2006; 281–283, DOI: 10.1101/pdb.prot4540.

Taher D, Solberg S, Prohens J et al. World vegetable center eggplant collection: Origin, composition, seed dissemination and utilization in breeding. Front Plant Sci 2017; 8: 1484, DOI: 10.3389/fpls.2017.01484.

Takagi H, Abe A, Yoshida K et al. QTL-seq: Rapid mapping of quantitative trait loci in rice by whole genome resequencing of DNA from two bulked populations. Plant J 2013; 74: 174–83, DOI: 10.1111/tpj.12105.

Tanksley SD, Ganal MW, Prince JP et al. High density molecular linkage maps of the tomato and potato genomes. Genetics 1992; 132: 1141–60, DOI: 10.1093/genetics/132.4.1141.

Tigchelaar EC, Janick J, Erickson HT. The genetics of anthocyanin coloration in eggplant (Solanum melongena L.). Genetics 1968; 60: 475, DOI: 10.1093/genetics/60.3.475.

Vilanova S, Alonso D, Gramazio P et al. SILEX: A fast and inexpensive high-quality DNA extraction method suitable for multiple sequencing platforms and recalcitrant plant species. Plant Methods 2020; 16: 1–11, DOI: 10.1186/s13007-020-00652-y.

Waterhouse AM, Procter JB, Martin DMA et al. Jalview Version 2-A multiple sequence alignment editor and analysis workbench. Bioinformatics 2009; 25: 1189–91, DOI: 10.1093/bioinformatics/btp033.

Wei Q, Wang J, Wang W et al. A high-quality chromosome-level genome assembly reveals genetics for important traits in eggplant. Hortic Res 2020; 7, DOI: 10.1038/s41438-020-00391-0.

Whitaker BD, Stommel JR. Distribution of hydroxycinnamic acid conjugates in fruit of commercial eggplant (Solanum melongena L.) cultivars. J Agric Food Chem 2003; 51: 3448–54, DOI: 10.1021/jf026250b.

Yasumura Y, Moylan EC, Langdale JA. A conserved transcription factor mediates nuclear control of organelle biogenesis in anciently diverged land plants. Plant Cell 2005; 17: 1894–907, DOI: 10.1105/tpc.105.033191.

