## Supplementary figures and images for "The irregular fruit green netting: An eggplant domestication trait controlled by the *SmGLK2* gene with implications in fruit colour diversification"

### Figure S1

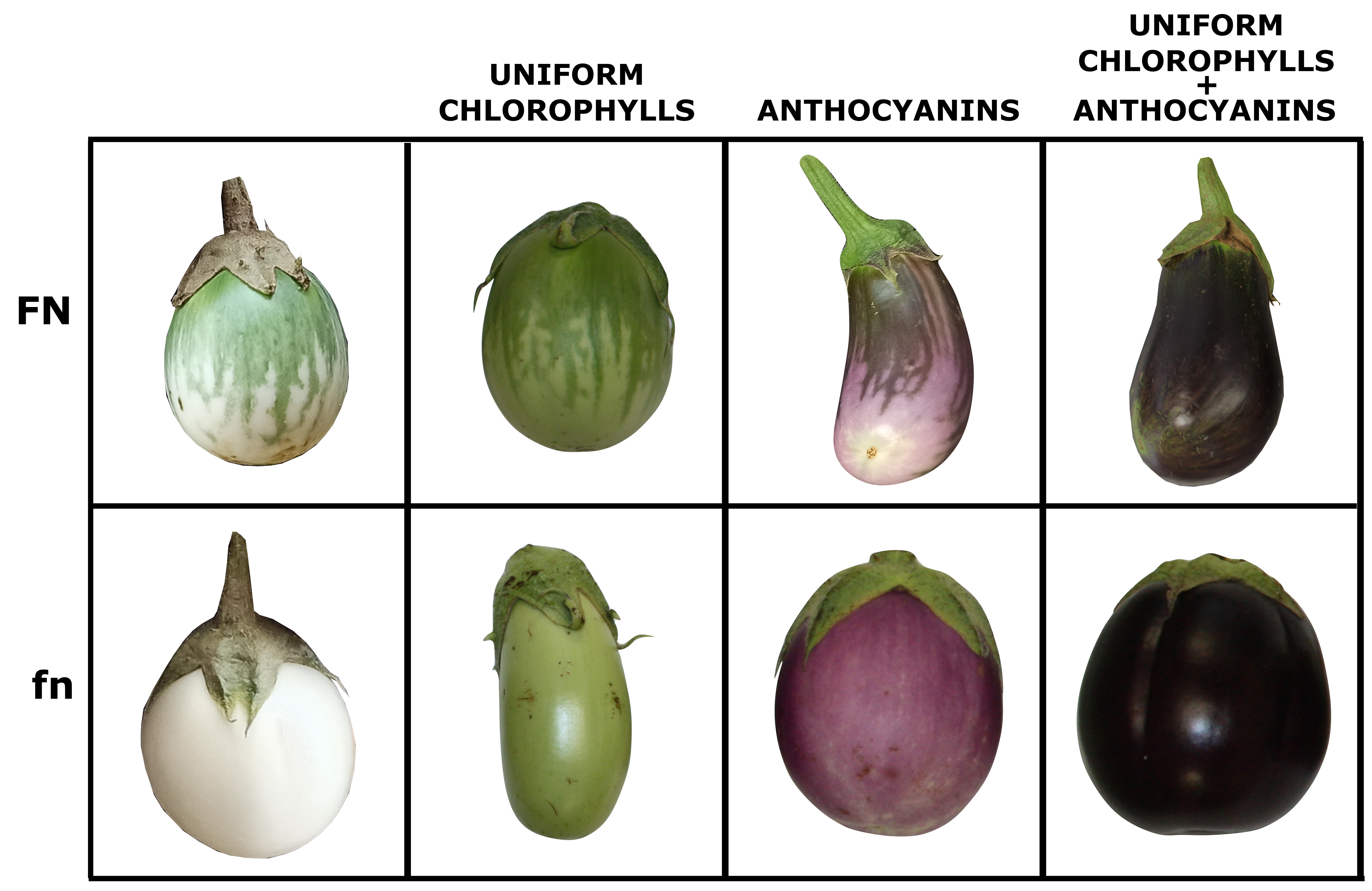
